# Dual origins for neural cells during development of the *Clytia* planula larva

**DOI:** 10.1101/2025.11.17.688882

**Authors:** Antonella Ruggiero, Anna Ferraioli, Sandra Chevalier, Pascal Lapébie, Romain Girard, Tsuyoshi Momose, Carine Barreau, Evelyn Houliston

## Abstract

Adult hydrozoan cnidarians undergo extensive tissue turnover, generating neural cell types including nematocytes (stinging cells) and gland cells from interstitial stem cells (i-cells) expressing stemness proteins such as Piwi and Nanos. The contribution of i-cells during embryogenesis, however, has been unclear. Here we address the origin of neural cells during development of the *Clytia hemisphaerica* ‘planula’ larva. Marker gene in situ hybridisation revealed that Piwi/Nanos1-expressing cells within the early gastrula presumptive endoderm generate a substantial pool of nematoblasts, a few of which migrate and differentiate in the planula ectoderm. Some neurogenic and neuronal markers, however, showed a markedly distinct expression profile, developing within a basal layer of the aboral/lateral ectoderm during gastrulation. Embryo bisection and lineage tracing experiments confirmed that sensory neurons and secretory cell types derive from gastrula ectoderm, while nematocytes and at least some ganglionic neurons derive from i-cells. Knockdown and inhibitor treatments revealed steps in neuron and nematocyte development regulated by Wnt-β-catenin. We conclude that two distinct neurogenesis pathways operate during *Clytia* embryogenesis, one involving aboral ectoderm delamination, and one generating mainly nematocytes from i-cell-like precursors.

**Summary statement:** During embryogenesis in the hydrozoan *Clytia* neural cell types derive both from *Piwi/Nanos* expressing “i-cells” and from ectodermal delamination during gastrulation.

## Introduction

Neural cells are one of the key animal innovations, and their developmental and evolutionary origins are thus much debated (Arendt et al., 2015; Burkhardt and Jékely, 2021; Hartenstein and Stollewerk, 2015; Hobert, 2025). Cnidarian species, including jellyfish and sea anemones, have a range of neural cell types and nervous systems, which have evolved in parallel to those of their sister clade Bilateria (Bosch et al., 2017; Kelava et al., 2015; Rentzsch et al., 2017).

These animals thus provide a valuable perspective on the underlying mechanisms of neurogenesis through comparisons with bilaterian model species. Current knowledge of neurogenesis in cnidarians is founded on extensive studies using adult *Hydra* polyps and during development of the sea anemone *Nematostella vectensis,* members respectively of the two main cnidarian clades Hydrozoa and Anthozoa. Hydra polyps undergo continuous cell turnover and propagate mainly by budding. The epithelia of their two body layers, ectoderm and gastroderm, renew themselves independently. Neural cell types, defined broadly to include ganglionic and sensory type neurons, stinging cells (nematocytes) and secretory gland cells, derive from a distinct population of multipotent stem cells called interstitial cells (i-cells), which also generate germ cells (Bosch et al., 2010; David, 2012; Frank et al., 2009; Siebert et al., 2019). I-cells are a specific feature of Hydrozoa. They typically have a rounded morphology and relatively large nucleus, and occupy spaces between the basolateral surfaces of the epitheliomuscular cells of the two adult body layers. They express orthologues of many genes associated with totipotency and germ cells, such as Nanos, Piwi, PL10 and Vasa (Genikhovich et al., 2006; Leclère et al., 2012; Rebscher et al., 2008). In the hydra polyp, neuron and nematocyte precursors derived from i-cells migrate into the ectoderm and gastroderm layers to take up their positions along the body axis and complete differentiation (Rentzsch et al., 2017). In another well-studied hydrozoan model, *Hydractinia*, single i-cells transplanted between polyps similarly give rise to neurons, nematocytes, gland cells and germ line, but unlike *Hydra* can also generate epithelial cells types (Varley et al., 2023). The i-cell to neuron, to nematocyte and to gland cell developmental trajectories in *Hydra* and *Hydractinia* polyps have been deduced from scRNAseq data (Cazet et al., 2023; Salamanca-Díaz et al., 2025; Siebert et al., 2019; Song et al., 2025) and tracked via expression of neurogenic transcription factors such as SoxBs (Chrysostomou et al., 2022). scRNAseq data have demonstrated equivalent trajectories in *Clytia* adult jellyfish (Chari et al., 2021).

While the i-cell derived pathways for neurogenesis in these adult hydrozoan polyps are well documented, the extent to which they also operate during initial embryonic development remains relatively poorly explored. Histological studies have described i-cells in the larval stage, the planula, largely confined to the endodermal (presumptive gastroderm) region (Bodo and Bouillon, 1968; Frank et al., 2009; Leclère et al., 2012), although whether they arise first in the endoderm or ectoderm may differ between species (see discussion in (Genikhovich et al., 2006)). Ganglionic-type and sensory-type neurons have been described in planulae, their neurites extending respectively from cell bodies close to the mesoglea or from the basal part of the cell, to form a network between the two tissue layers of the larva (Attenborough et al., 2019; Martin, 2000; Piraino et al., 2011; Ramon-Mateu et al., 2025). Martin and Thomas provided experimental evidence from studies in hydrozoan planulae from various species that ganglionic cells derive from i-cells in the endoderm (Martin, 1988; Martin, 1991; Martin and Thomas, 1980; Martin and Thomas, 1981; Thomas et al., 1987). These experimental studies also indicated, however, that at least some planula neural cell types derive directly from the ectodermal layer. Notably, neurosensory, neurosecretory (“cellule glandulaire sphéruleuse”) and mucous cells (“cellule glandulaire spumeuse”) were identified by electron microscopy in planulae derived from aboral halves of *Clytia* (*Phialidium) gregarium* gastrula-stage embryos, which lack i-cells as well as presumptive gastroderm (Thomas et al., 1987). In contrast, nematocytes and ganglionic cells were found only in larvae derived from oral halves.

Similarly, when i-cells were depleted from developing larvae from the hydrozoan polyp species *Pennaria* by colchicine treatment, planulae were preferentially depleted in nematocytes and ganglionic cells (Martin and Thomas, 1981).

Unlike hydrozoans, anthozoan cnidarians do not have a distinct i-cell population, although recent studies in *Nematostella* have identified cells with similar characteristics in the gastrodermal mesenteries of the adult (Denner et al., 2024; Miramón-Puértolas et al., 2024), and shown that neuronal and gland cells have common progenitors (Steger et al., 2022).

During embryo development, neural precursors arise in a salt and pepper fashion within the two epithelial body layers. A first wave of neurogenesis during gastrulation generates sensory and ganglionic neurons in the ectoderm, and in a second wave, during the planula stage, precursors arise similarly in the gastroderm (Kelava et al., 2015; Lemaître et al., 2023; Nakanishi et al., 2012; Rentzsch et al., 2017; Richards and Rentzsch, 2014; Richards and Rentzsch, 2015).

We have addressed the origin of neural cell types during larval development in *Clytia hemisphaerica.* Distinct cell types in the *Clytia* planula larvae that can be together considered neural on the basis of shared transcriptomic features include sensory neurons, neurosecretory cells and mucous cells types interspersed in the ectoderm layer, as well as ganglionic-type neurons with cell bodies close to the mesoglea, and nematocytes (Ferraioli et al., 2026). In *Clytia* species, gastrulation proceeds by ingression of presumptive gastroderm cells from the oral pole (Byrum, 2001), carrying with them the i-cells (Leclère et al., 2012). We first mapped nematogenesis and neurogenesis by in situ hybridisation using molecular markers. We then tested the contribution of the ectoderm to neurogenesis by gastrula bisection and lineage tracing experiments. Finally we assessed the influence of Wnt-β-catenin signalling on neural development using antisense Morpholino oligonucleotides targeting Wnt3 (Wnt3-MO; Momose et al., 2008) and timed treatments with the pharmacological Wnt-β-catenin pathway inhibitor PKF118-310 (Lepourcelet et al., 2004). Our findings demonstrate that the neural cell types of the *Clytia* planula larva develop during gastrulation by two parallel pathways, one generating sensory neurons along with secretory cells types from a basal layer of aboral/lateral ectoderm, and the other operating within the gastrodermal region to mainly produce nematocytes from i-cell like precursors.

## Materials and Methods

### Manipulation of Eggs and Embryos

Eggs and embryos were obtained from laboratory-raised Z strain medusae (Leclère et al., 2019) and cultured in 0.2 µm Millipore filtered ASW (Red Sea Salt brand; (Lechable et al., 2020). Medusae were kept on daily dark-light cycles with gametes collected two hours after light. Blastula bisection and blastomere separation were performed using fine Tungsten wire loops in agar-lined petri dishes as described previously (Leclère et al., 2012). PKF118-310, which interferes with the interaction between β-catenin and the conserved transcription factor Tcf/LEF required to regulate gene expression via this pathway (Lepourcelet et al., 2004), was used at 0.8 µM in FASW, diluted from a 15 mM stock immediately before use.

### Dendra2 lineage tracing

The coding sequence for Dendra2 (Chudakov et al., 2007) was adapted to the codon usage of *Clytia* (Weissbourd, 2021). A T3 promoter-Dendra2 ORF DNA template was obtained by PCR and mRNA was synthesised using mMessage mMachine^TM^ T3 Transcription Kit (Ambion), followed by poly-A addition using the Poly(A) Tailing Kit (Ambion). 0.4 - 1.0µg/µl mRNA was injected into freshly spawned eggs prior to fertilisation using an Eppendorf nanoject injection system as described in detail previously (Uveira et al., 2025), aiming at 3-5% egg volume. Embryos were allowed to develop at 18°C until the 4 or 8 cell stage then cultured 16.5°C overnight (approx 18 hours total) until the early gastrula stage. They were then temporarily mounted between slides and coverslips separated by plasticine spacers compressed to immobilise nearly completely the larvae during photoconversion, performed on a Leica Stellaris confocal microscope using the LasX software FRAP mode with a 10X objective. Each defined region, either oral or aboral, was exposed to 30 seconds scanning of the 405 nm Laser beam, at 100% power. Embryos were then cultured at 17°C in MFSW until planula stage before fixation for immunofluorescence staining.

### Fluorescence microscopy

For immunofluorescence, samples were fixed for 2 hours at room temperature in IF fix [0.1 M Hepes (pH 6.9), 50 mM EGTA (pH 7.2), 10 mM MgSO4, 80 mM maltose, 0.2% Triton X-100, 4% paraformaldehyde] before proceeding to three washes in Phosphate Buffered Saline (PBS; 110 mM NaCl, 1.9 mM KCl, 8 mM Na_2_HPO_4_, 2 mM KH_2_PO_4_ pH 7.2) containing 0.01% Triton X-100 for 10 min or longer, extraction in PBS-0.2% Triton for 10 minutes, two washes in PBS-0.01% Triton X-100, and one rinse in PBS. Primary antibodies were an affinity purified rabbit polyclonal antibody raised against the peptide GRFamide (gift of Gáspár Jékely) or anti FMRFamide (Immunostar Product ID: 20091) and an anti-LWamide mouse monoclonal antibody (gift of O. Koizumi (Koizumi et al., 2015)). Samples were incubated overnight at 4°C with primary antibodies and/or Alexa488- or Rhodamine-Phalloidin (1:100 from stocks in methanol; Fisher Scientific) to visualise the actin-rich cell cortices, followed by washes in PBS-0.01% Triton X-100 (three times for 10 min or longer) and then incubation in rhodamine, Alexa555 or Alexa647 coupled anti-rabbit or anti-mouse IgGs (Jackson ImmunoResearch, dilution 1:100) for 3 hours at room temperature, with Hoechst dye 33342 (Bisbenzimide from Sigma; 0.3 μg/ml final) included to stain DNA. After washes in 0.01% PBS-Triton X-100 (three times each for 10 min or longer) and a PBS rinse, specimens were mounted in Citifluor mounting medium (Citifluor Ltd) and imaged using Leica SP5, SP8 or Stellaris confocal microscopes.

### Gene cloning

In situ hybridisation probes were synthesised from cDNA clones retrieved from our EST collection when available (Chevalier et al., 2006). For the remaining sequences, cDNAs were cloned by PCR using the pGEM-T Easy system (Promega). Accession numbers for the newly identified sequences, and XLOC identifiers (Chari et al., 2021; Leclère et al., 2019) for all genes analysed, are provided in Supplementary File S1.

### In situ hybridisation

DIG-labeled and fluorescein-labeled antisense RNA probes for in situ hybridisation were synthesised using Promega T3/T7/Sp6 RNA polymerases, purified using ProbeQuant G-50 MicroColumns (GE Healthcare) and stored at -20°C in 50% formamide. Planulae were fixed for in situ hybridisation in a cold solution of 3.7% formaldehyde and 0.2% glutaraldehyde in PBS on ice for 40 minutes to 2 hours, fixation time depending on the developmental stage. Specimens were then washed with PBST (1X PBS + 0.1% Tween 20), dehydrated stepwise in methanol on ice, and stored in 100% methanol at -20°C. Samples were fixed after culture for 24, 48 or 72 hours post fertilisation (hpf) at 17–18°C.

In situ hybridisation was performed either using the InsituPro robot (Intavis®) as described previously (Lapébie et al., 2014, modified as in Chari et al., 2021) or by hand as described in Sinigaglia et al. (2020).

## Results

### Nematoblast production from i-cells during gastrulation

We tracked the formation of neurons and nematocytes during *Clytia* embryogenesis using in situ hybridisation (Figure 1). We targeted genes identified previously as expressed in i-cells and/or during neurogenesis/nematogenesis by trajectory inference analysis of scRNAseq data from *Clytia* adult medusa (Chari et al., 2021). We included *Piwi* and *Nanos1* probes as markers of i-cells (Leclère et al., 2012). Note that this operational identification of ‘i-cells’ used here includes stem cells along with cells that have already embarked on developmental pathways to form neurons and nematocytes, as shown by the onset of lineage-specific gene expression (Chari et al., 2021; Denker et al., 2008; Mochizuki et al., 2000; Siebert et al., 2008; Varley et al., 2023). As found previously (Leclère et al., 2012), cells expressing high levels of Nanos1, Piwi and Vasa were detected in the oral pole region at the early gastrula stage, then concentrated within the oral part of the presumptive gastroderm region at the end of gastrulation at the parenchymula (24 hours after fertilisation: P1) stage, and finally spread within this region along the oral-aboral axis of fully formed planulae (second and third day following fertilisation : P2 and P3 stages; Fig. 1A).

**Figure 1.**
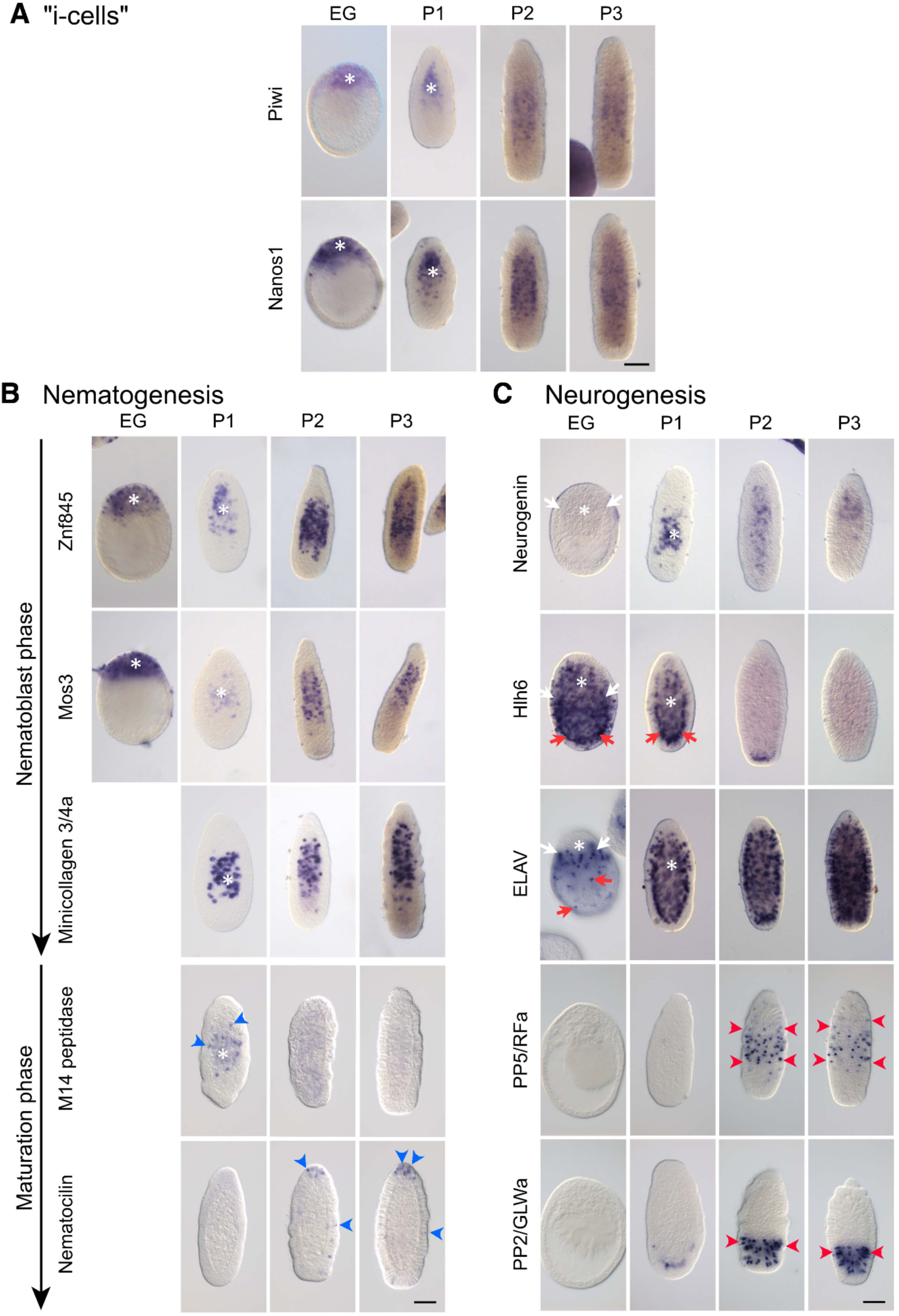
Expression of nematogenesis and neurogenesis genes during planula development. In situ hybridisation images of embryos fixed at early gastrula (EG), parenchymula (P1; corresponding to the end of gastrulation; see Kraus et al., 2020), 2-day (P2) and 3-day (P3) settlement-competent planula larvae, detecting the following mRNAs: (A) Piwi and Nanos1 as markers of “i-cells” (which under this operational definition includes also early stages of derivative cell types); (B) Znf845, Mos3 and Minicollagen 3/4a expressed during the first phase ‘nematoblast formation’ as indicated by the top arrow, and M14 peptidase and Nematocilin during the second phase (nematocyte maturation; bottom arrow); (C) Transcription factors Neurogenin and Hlh6, and neuronal markers ELAV, PP5 (RFamide precursor) and PP2 (GLWamide precursor). Oral pole is at the top in all panels. White asterisks indicate ingressing presumptive gastroderm and white arrows the extent of its ingression. Blue arrowheads indicate differentiating nematocytes migrating from gastroderm to ectoderm layers during the maturation phase of nematogenesis. Red arrows show cells expressing Hlh6 and ELAV lining the blastocoel during gastrulation and in a layer between ectoderm and gastroderm layers at P1. Red arrowheads indicate the neuronal cell bodies in the planula ectoderm. Scale bars: 100 µm.

The formation of stinging cells, or nematogenesis, can be separated into two distinct phases as characterised from *Clytia* medusa scRNAseq data (Chari et al., 2021). A ‘nematoblast’ phase involving construction of the nematocyst capsule is followed by a ‘maturation’ phase during which a functional nematocil is assembled. These phases have almost non-overlapping transcriptome signatures, so can be distinguished by in situ hybridisation using specific markers (Fig. 1B). Amongst the earliest expressed genes in the transition from i-cells to nematoblasts, as revealed from analysis of medusa scRNAseq data, are *Znf845* and then *Mos3* (Chari et al., 2021). We detected transcripts for these two genes strongly at the oral pole of the early gastrula embryo (Fig. 1B), consistent with their original identification as orally-expressed early zygotic genes (Lapébie et al., 2014). Cells expressing these genes moved into the blastocoel along with the mass of mesenchymal presumptive gastrodermal cells, their expression patterns until the end of gastrulation mirroring those of *Piwi* and *Nanos1*. In planula stages, expression of all these genes continue to show a similar distribution, although the expression of *Znf845* and *Mos3* becomes progressively more pronounced. Together these observations suggest that many of the *Piwi*/*Nanos1* expressing cells detected during planula development are early stage nematoblasts rather than stem cells.

Expression of *Minicollagen 3/4a* mRNA, which codes for a structural component of the nematocyst capsule (Denker et al., 2008) expressed during nematoblast formation but strongly downregulated during the subsequent phase of nematocyte maturation and migration (Chari et al., 2021; Condamine et al., 2019), was detected from the parenchymula (P1) stage. In P2 and P3 planulae, nematoblasts are positioned towards the basal side of the developing gastroderm epithelium, occupying a central “trunk” position along the oral-aboral axis (Fig. 1B; Bodo and Bouillon, 1968).

Despite the presence of abundant nematoblasts in P2 and P3 planula larvae, we detected relatively few cells undergoing nematocyte maturation. mRNA for the M14 peptidase gene, whose expression marks the transition from nematoblast to terminal differentiation, could be detected in some central cells within the gastroderm at P2. We interpret these as a sub-population of the nematoblasts starting to differentiate at this stage. Fewer such cells were observed at P3 suggesting that differentiation of a small population of nematoblasts occurs as a wave. Reciprocally, mRNA for Nematocilin, a component of the nematocil trigger structure and a marker of mature nematocytes, became clearly detectable by P3 in the ectodermal layer, scattered along the trunk, and clustered at the oral pole (Fig. 1B). These expression patterns confirm that nematocyte differentiation accompanies migration into the ectoderm, whilst indicating also that only a small subpopulation of nematoblasts continue development to form mature nematocytes prior to larval metamorphosis.

Overall these profiles of nematogenesis gene expression indicate i) that nematoblasts start to develop at the onset of gastrulation as an abundant Piwi/Nanos positive cell population in the presumptive gastroderm at the oral pole, and ii) that development of most of this nematocyte pool becomes arrested at the end of the nematoblast formation phase.

### Mixed expression profiles for neurogenic and neuronal markers

We similarly tracked by in situ hybridisation the development of neurons (Fig. 1C). We included probes for two transcription factors expressed along the i-cell to neuron trajectory in the scRNAseq analyses from *Clytia* medusae (Chari et al., 2021): the atonal family Hlh6, orthologue to *Nematostella* Ath-like (Richards and Rentzsch, 2014), and Neurogenin. These two transcription factors showed very distinct expression profiles (Fig. 1C). Neurogenin mRNA could be detected in a few scattered cells within the endoderm from 24h, consistent with the development of neuronal precursors from i-cells. In contrast, Hlh6 showed strong expression much earlier, at the gastrula stage, in cells lining the blastocoel, covering lateral and aboral regions. The expression profile of Hlh6 is reminiscent of that of the SoxB2 family transcription factor Sox10 (Kraus et al., 2020; expression profiles at early gastrula stages compared in Fig. 2A, B). Like Sox10, Hlh6 expression diminishes after gastrulation. These observations suggest that a layer of cells with neurogenic potential is present transiently during late blastula and gastrula stages. Correspondingly, expression of an ELAV gene expressed in most planula neurons (identifier XLOC_030971; Ramon-Mateu et al, 2025) was detected in cells lining the blastocoel during gastrulation, as well as in cells within the ingressing presumptive gastroderm potentially associated with i-cells (Fig. 1C and Fig. 2C). At the beginning of gastrulation, ELAV positive cells were detected scattered across the inner surface of the blastocoel, stronger expression observed closest to the oral pole (Fig. 2C). As gastrulation progressed, ELAV expressing cells accumulated in a corresponding layer between the ingressing gastroderm and the ectoderm, as well as within both tissue layers.

**Figure 2.**
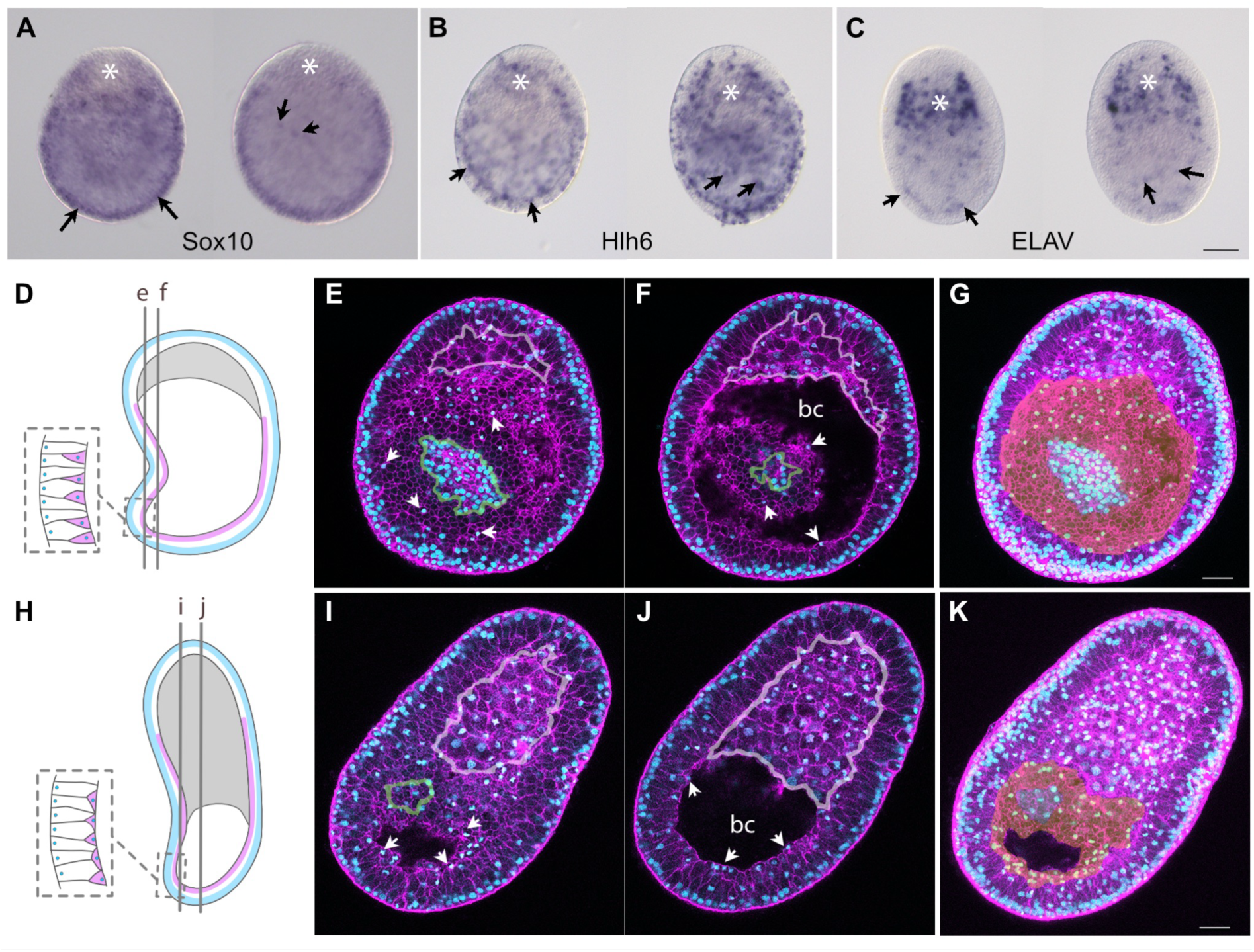
Formation of an inner ectodermal layer during gastrulation. (A-C) In situ hybridisation images with probes for Sox10 (A), Hlh6 (B) and ELAV(C) of early gastrula embryos showing focal planes through the centre of the blastocoel (left) and basal level of the ectoderm (right) for each probe. Asterisks mark the centre of the invaginating presumptive gastroderm. Arrows show examples of stained cells in the ectoderm. (D-K) Confocal analysis to demonstrate the developing inner ectodermal layer. Early (D-G) and mid (H-K) gastrula stage embryos, illustrating stratification of the presumptive ectoderm. Presumptive gastroderm cells are ingressing from the oral pole (top right in both embryos). (D and H) Cartoons of the two embryos indicating the planes of confocal sections shown in E, F, I and J. Undulations are present after fixation and mounting. The position of the putative neurogenic layer is shown in purple, and of the apical nuclei of the ectodermal cells in blue. Drawings in the dashed boxes indicate stratified cellular organisation of the corresponding areas; the purple basal cells may retain thin apical projections reaching the embryo outer surface (‘pseudo-stratification’). (E, F, I and J) Confocal images each from z stacks. Actin-rich cell cortices were stained with rhodamine-phalloidin (magenta) and DNA with Hoechst 33342 (cyan). Distance between planes e and f was 14.8 µm, between i and j was 32.2 µm. Arrowheads indicate some examples of the nuclei of putative delaminating cells. bc=blastocoel. Ingressing presumptive endoderm cells are outlined in pink. Apical nuclei of the ectoderm visible due to folding are outlined in green. (G and K) Maximum projection images of the same z stacks (5 planes over a total thickness of 9.6 µm for G; 5 planes over a total thickness of 32.2 µm for K). These projections highlight the layer of nuclei lining the blastocoel, coloured with an orange overlay. Scale bars 25 µm.

These expression patterns suggest that the formation of at least some planula neural cell types occurs by stratification of the ectoderm during late blastula and gastrula stages, involving sequential expression of Sox10, Hlh6 and then ELAV in the basal layer. Stratification (or pseudo-stratification) was previously suggested by our study of cell morphogenesis during *Clytia* gastrulation based on confocal and electron microscopy (Kraus et al., 2020). Nuclei were found to appear progressively on the basal side of the ectoderm to line the blastocoel during gastrulation, in parallel with the EMT (Epithelio Mesenchymal Transition)-based ingression of presumptive gastroderm cells from the oral pole. We re-examined confocal z-stacks of gastrula stage embryos stained to highlight cell boundaries and nuclei (Figure 2), constructing maximum-projection images to visualise the emerging layer of cells (coloured orange in Fig. 2G and 2K). This layer mirrors the distribution of Hlh6/ELAV expressing cells at these stages ( Fig. 2), supporting the idea that it is neurogenic. We cannot tell from this analysis whether the stratification involves complete or incomplete cell delamination, as illustrated in Fig. 2D and 2H).

To detect differentiated neurons at planula stages we monitored expression of the RFamide neuropeptide precursor PP5 and the GLWamide precursor PP2. RFamides are widely expressed in cnidarian neurons of various types including ganglionic type cells and in putative planula sensory cells in some species (Attenborough et al., 2019; Hayakawa et al., 2007; Piraino et al., 2011; Weissbourd, 2021). GLWamides, involved in mediating the larval settlement response in many species of cnidarian (Takahashi and Hatta, 2011), have been detected in sensory cells distributed at or around the aboral pole in *Hydractinia* and *Clytia*, and in a more widely distributed ganglionic cell population in *Clava* (Gajewski et al., 1998; Piraino et al., 2011; Ramon-Mateu et al., 2025). During development we first detected neurosensory cells expressing the GLWamide precursor PP2 (Ramon-Mateu et al., 2025) at the parenchymula stage, and then more strongly in P2 and P3 planulae, in the aboral ectoderm (Fig. 1C). Cells expressing the RFamide precursor PP5, likely ganglionic neurons, were detected in P2 and P3 planulae in cell bodies in the trunk region of the ectoderm close to the mesoglea (Fig. 1C). These two neuropeptide precursors in *Clytia* thus allow subpopulations of neurons positioned differentially along the body axis with aboral secretory versus ganglionic morphologies to be distinguished.

### Ectodermal neurogenesis demonstrated by gastrula bisection experiments

To test whether neural cell types derive from the gastrula ectoderm during *Clytia* gastrulation we used two approaches. Firstly, we bisected early-mid gastrula stage embryos and fixed them at the planula stages to assess neural cell development by fluorescence microscopy or in situ hybridisation (Figure 3).

**Figure 3.**
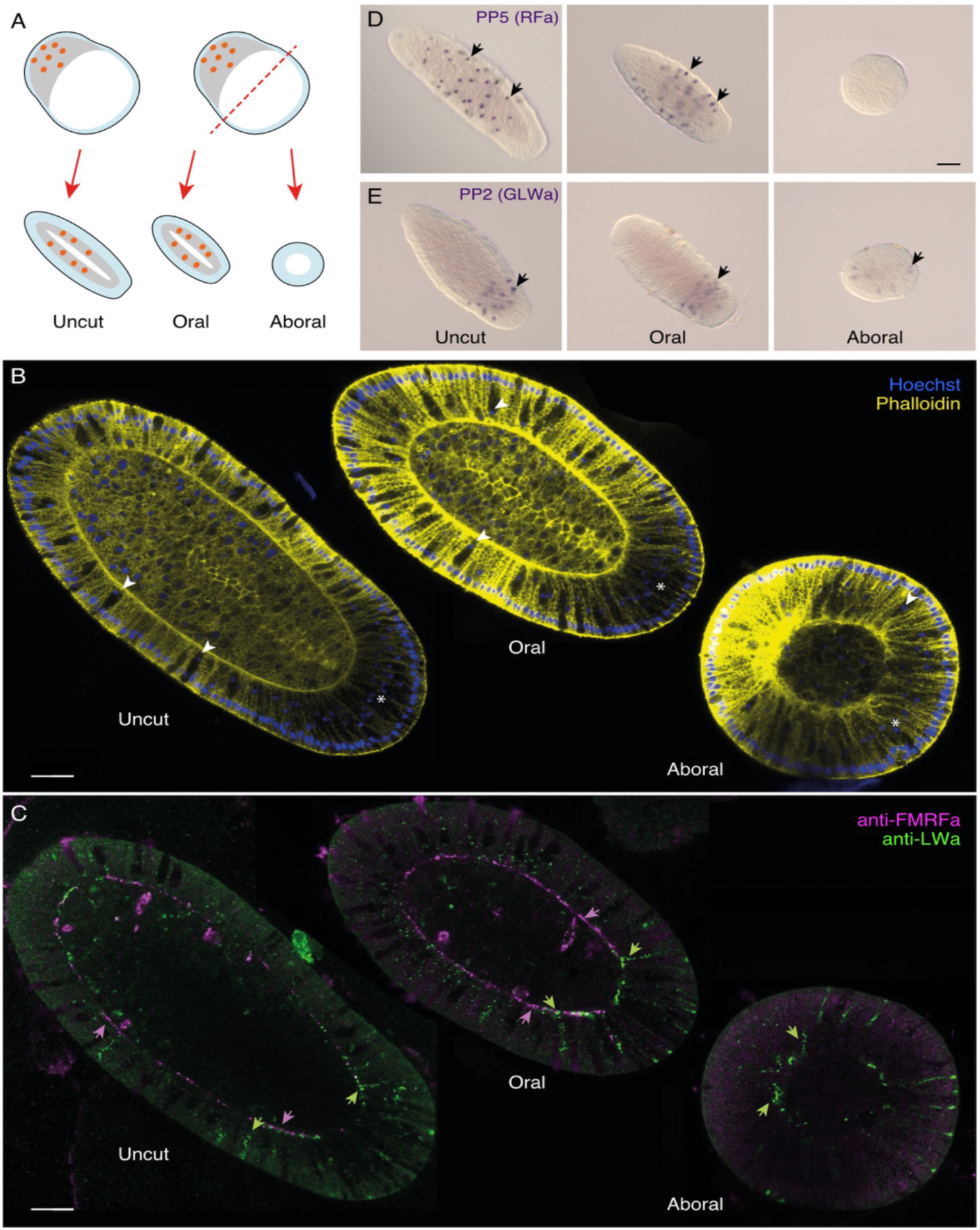
Gastrula bisection demonstrates ectodermal origin for some neural cell types. (A) Cartoon of experimental design. Unmanipulated early-mid gastrulae were allowed to develop to planula stages (48-66 hpf). Other embryos were bisected into oral and aboral parts and then allowed to develop in parallel. Orange dots indicate i-cells positioned in the ingressing presumptive endoderm region (grey) at the oral pole. (B) Overlaid confocal images of Hoechst staining of nuclei (blue) and Alexa488-Phalloidin staining (yellow). Endogenous GFP expressed in epitheliomuscular cells is also visible in this channel (Fourrage et al., 2014). Uncut, Oral and Aboral derived planulae from left to right, all fixed on the second day after fertilisation. White arrows show examples of mucous cells; white asterisks mark the zone of aboral secretory cells, which have medially positioned nuclei and do not express GFP. (C) Confocal images of the same planulae as in (B) stained with anti-FMRFamide (magenta) and anti-LWamide (green). Examples of neurites stained with each antibody are shown with magenta and green arrowheads respectively. (D-E) In situ hybridisation to detect PP5 and PP2 expression in planula fixed at 48 hpf. Arrows show examples of stained cells. Uncut, Oral and Aboral derived planulae from left to right. Oral is top left in all panels. Scale bars 50 µm.

Phalloidin staining confirmed that aboral-derived planulae lacked gastroderm, and also showed that the ectoderm contained differentiated mucous cells and aboral secretory cells (Fig. 3B). These can be distinguished by their morphology, lack of endogenous GFP expression, and nuclei positioned at basal or medial positions within the ectoderm, in contrast to the surrounding GFP-rich epitheliomuscular cells with apical nuclei. Anti-FMRFamide antibodies, which recognise RFamide and related ganglionic cells, and anti-LWamide antibodies recognising aboral sensory cells, both detected distinct populations of ganglionic and sensory type neural cells in equivalent distributions in all uncut planulae and planulae derived from oral halves (16/16 and 22/22 planulae examined respectively from 2 experiments). In contrast, anti-LWamide positive neurons but no FMRFamide neurons were detected in most planulae derived from aboral halves (Fig. 3C; Of 26 double-stained planulae examined from 2 experiments, 26 showed LWamide positive and 1 showed FMRFamide positive cells). Equivalent results were obtained in two in situ hybridisation experiments with probes detecting precursor mRNAs for RFamide (PP5; Fig. 3D) and GLWamide (PP2; Fig. 3E). In the first experiment 28/28 oral-derived planulae and 1/11 aboral-derived planulae showed PP5 positive cells, while 2/2 oral-derived planulae and 7/7 aboral-derived planulae showed PP2 positive cells (Fig. 3D-E; fixations at 48 hpf). In a second experiment (fixations at 66 hpf), 14/14 oral-derived planulae and 3/11 aboral-derived planulae showed PP5 positive cells, while 23/25 oral-derived planulae and 11/12 aboral-derived planulae showed PP2 positive cells.

These embryo bisection experiments thus demonstrate that mucous cells, secretory cells and GLWamide sensory neurons can form from early gastrula ectoderm. They also are consistent with the conclusion that ganglionic neurons including RFamide neurons originate from the oral pole of the gastrula, and specifically from i-cells, although the absence of an oral Wnt signalling centre in aboral-derived embryos could account, at least in part, for the lack of detectable PP5/RFamide expression, an idea developed further below.

### Dendra2 tracing supports dual origins for neural cells

To confirm the ectodermal origin of GLWamide neurons and test the origin of RFamide neurons we performed lineage tracing using the photoconvertible fluorescent protein Dendra2 (Chudakov et al., 2007). We injected mRNA coding for Dendra2 into eggs prior to fertilisation and photoconverted the protein from green to red in patches of cells at either the aboral or oral poles of the embryo at early-mid gastrula stages using the 405nm laser on a confocal microscope. The embryos were then cultured to the planula stage (P2) and the identity of Dendra2 red positive cells analysed. As illustrated in Fig. 4A, we reason that in cases of aboral photoconversion the target zone in the early gastrula does not include presumptive gastroderm or i-cells, implying that any Dendra2 red cells detected at planula stages must derive from the aboral ectoderm. Conversely, oral Dendra2 photoconversion zones at early gastrula stages are predicted to include part of the presumptive oral ectoderm and gastroderm, and some of the i-cells. During gastrulation these will ingress aborally with the developing gastroderm to occupy the central region of the developing planula, before some of their neuron and nematocyte descendants migrate out into the ectoderm. We thus anticipate finding Dendra2-red positive nematocytes and neurons in the ectoderm at planula stages, both within the oral Dendra2-red positive patch, but also individually in more aboral positions separated spatially from this patch.

**Figure 4.**
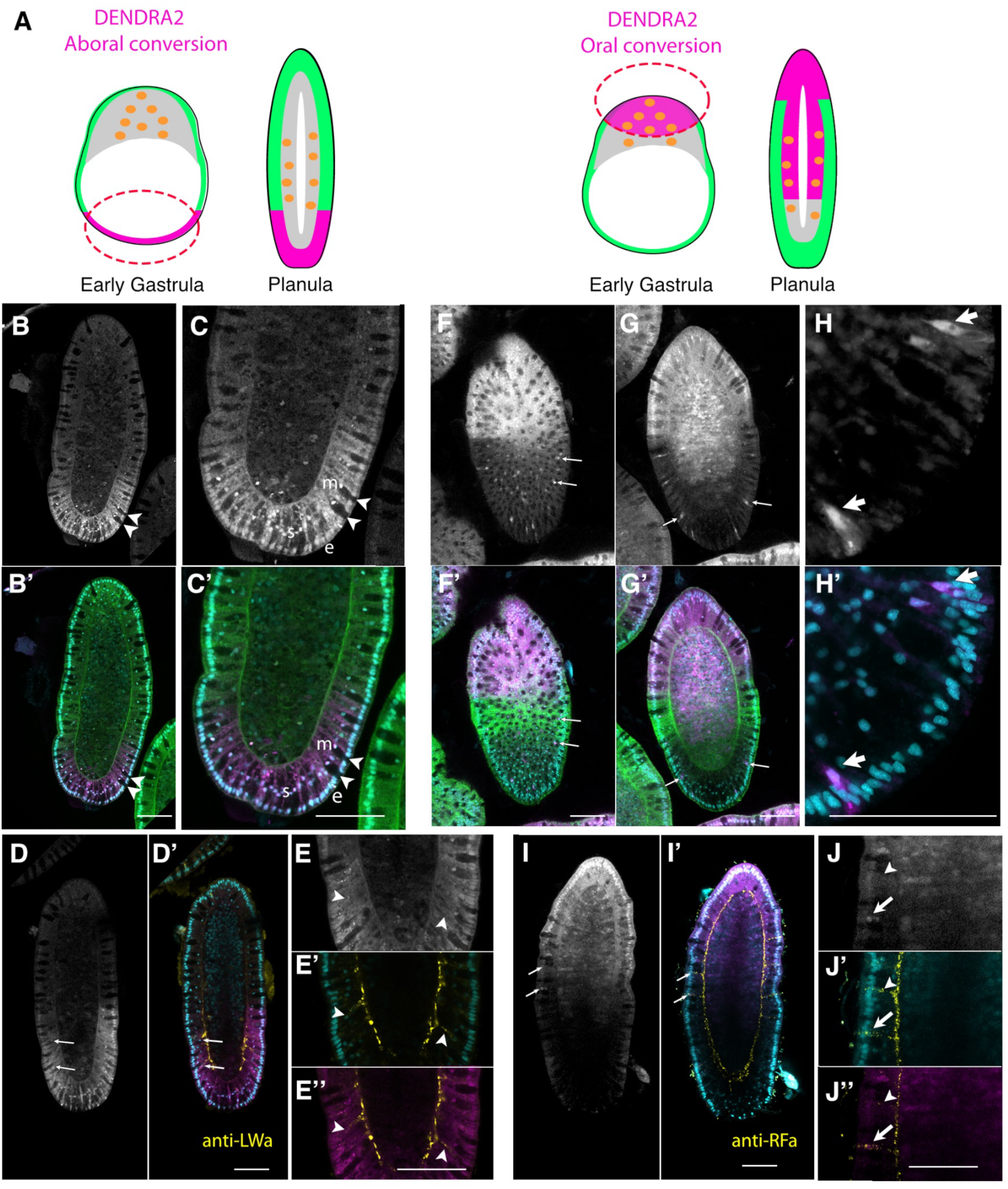
Dendra2 photoconversion traces neural fates from the gastrula stage. (A) Experimental design; Regions of the aboral or oral poles of early gastrula stage embryos expressing Dendra2 from mRNA injected into the egg were photoconverted to red using the 405nm laser of a Leica Stellaris confocal microscope. (B-J) Confocal microscope images of planulae derived from either aboral (B-E) or oral (F-J) conversions. B, C, D, E, F, G, H, I and J show the distribution of cells containing Dendra2-red. Other panels show the same planes with the Dendra2-red image in magenta overlaid with Phalloidin-Alexa488/endogenous GFP (green) and Hoechst staining of nuclei (cyan) in B’, C’, F’ and G’; with Hoechst (cyan) only in H’; with anti-LWamide staining (yellow) and Hoechst (cyan) in D’; with anti-LWamide staining (yellow) only in E’’; with anti-RFamide staining (yellow) and Hoechst (cyan) in I’; with anti-RFamide staining (yellow) only in J’’. Panels E’ and J’ show the overlay of Hoechst staining (cyan) with anti-LWamide staining or anti-RFamide staining (yellow) respectively. (B-C) Examples of aborally derived, cytoplasm-poor, mucous cells appearing black with basal nuclei (arrowheads). Nuclei in the mucous cells (m), aboral secretory cells (s) and epithelial cells (e) are all positive for Dendra2-red. (D-E) Examples of LWamide positive cells in the aboral ectoderm (arrows and arrowheads) positive for Dendra2-red. (F-H) Examples of nematocytes (arrows and arrowheads) in ectodermal regions far from the oral-irradiated epithelial clone, likely derived from gastrodermal i-cells. (I-J) Examples of RFamide positive cells in the trunk ectoderm (arrows and arrowheads). The lower cell (arrow) is positive for Dendra2-red, so is deduced to derive from a photoactivated cell in the involuting presumptive gastroderm region; the upper cell (arrowhead) is negative so could derive from a non-photoactivated presumptive endoderm cell or an ectoderm cell. Oral pole is at the top in all panels. Each image is through a central plane of the planula except F/ F’ which is closer to the surface, within the ectoderm layer. Scale bars all 50 µm.

As expected, the progeny of aboral photoconverted Dendra2 cells formed coherent aboral patches of aboral ectoderm by the planula stage, while the progeny of oral photoconverted cells included oral ectoderm and gastroderm, the exact time and zone of photoconversion in each embryo determining the precise pattern observed (Fig. 4B-J). Confirming the results of the gastrula bisection experiments, the patches of photoconverted Dendra2 progeny in planulae derived from aboral (ie ectoderm-restricted) irradiation included mucous and secretory cells, characterised by their typical morphologies and nuclei positioned respectively in basal or medial positions (Fig. 4B-C). In addition, some Dendra2-red cells in the aboral ectoderm were stained by the anti-LWamide antibody (Fig. 4D-E).

In planulae derived from oral photoconversion, Dendra2-red nematocytes were clearly detected in the ectoderm far from the oral patch of photoconverted epidermal cells, validating this technique as an indicator of i-cell origins (Fig. 4F-H). Furthermore we could detect RFamide in Dendra2 red cells at positions detached from the oral ectoderm patch, but not scattered as widely as the nematocytes (Fig. 4I-J).

We conclude from these two approaches that during *Clytia* planula development from the gastrula stage, neural cell types have two origins. I-cells in the gastrodermal region provide nematocytes and at least some of the RFamide ganglionic cells, whereas GLWamide sensory cells, along with mucous and aboral secretory cells, derive mainly from gastrula ectoderm.

### Wnt-β-catenin signalling promotes nematogenesis and neurogenesis

Wnt-β-catenin signalling has widespread roles in regulating neurogenesis across animals as well as body axis specification. In *Clyta* it determines the oral-aboral axis, with a key role for the ligand Wnt3, expressed orally (Momose et al., 2008; Uveira et al., 2025), while in *Hydractinia* it also regulates i-cell differentiation into neurons (Duffy et al., 2010; Duffy et al., 2012). Conversely, pharmacological stimulation of the Wnt-β-catenin pathway in *Hydractinia* and *Hydra* polyps causes excess formation of nematocytes and of some nerve cell types (Hensel et al., 2014; Khalturin et al., 2007; Teo et al., 2006). In *Hydra* it has been proposed that Wnt activity promotes nematocyte differentiation for the mouth and tentacle structures at the oral tip of the polyp where this pathway is active (Bosch and David, 1991; Hobmayer et al., 2000; Khalturin et al., 2007; Lengfeld et al., 2009).

To test the role of Wnt-β-catenin signalling in *Clytia* nematogenesis/neurogenesis, we used two approaches: Wnt3-MO injection, prior to fertilisation, which prevents oral territory specification and delays gastrulation (Lapébie et al., 2014; Momose et al., 2008), or timed incubations in PKF118-310, which interferes with TCF/β-catenin interactions (Chudekov et al., 2007; Figure 5). Early PKF118-310 treatments (from cleavage stage to 24 hpf) have been shown to mimic the morphological aboralising effects of Wnt-β-catenin signalling inhibition on axis establishment (Sinigaglia et al., 2020), which is completed by the parenchymula stage; Late treatments (24 to 48 hpf) were used to specifically target the period of neuronal differentiation. These late treatments did not shift the established expression territories for Wnt3, WegA1 or FoxQ2a, but did abolish Bra1 expression at the oral tip of the planula (Supplementary File S2), indicating that once established axial identities along the oral-aboral axis can be largely maintained independent of Wnt-β-catenin signalling. We monitored development of nematocytes and neurons in these experiments using the panel of in situ hybridisation probes presented in Figure 1. The staining was assessed for each group by counting the embryos/larvae showing similar numbers of stained cells as untreated embryos (scored as “positive”), either indistinct staining or of just a few cells (scored as “trace”), or no cellular staining (scored as “none”) (Supplementary File S3).

**Figure 5.**
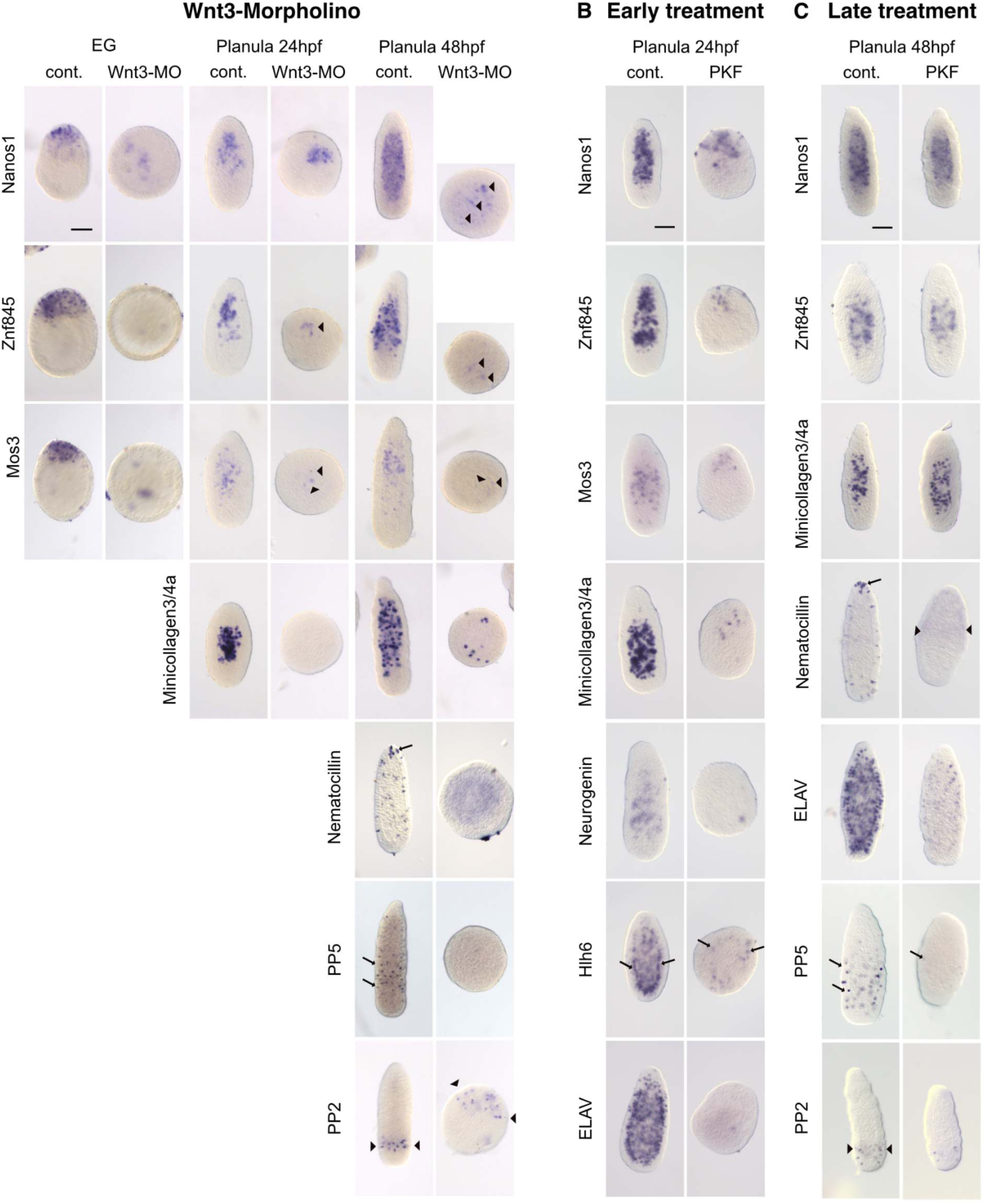
Regulation of nematogenesis and neurogenesis by Wnt-β-catenin signalling. In situ hybridisation images of embryos fixed at the early gastrula (EG); 24 hpf parenchymula (P1) and 48 hpf planula larvae (P2) stage with probes to detect the genes indicated to the left of each panel. Majority phenotypes are shown (see numbers in Supplementary File S3). (A) Uninjected embryos (cont.) compared with Wnt3-Morpholino injected embryos. (B) Untreated embryos (cont.) compared with embryos incubated from 4 to 24 hpf in PKF118-310 (PKF). (C) Untreated embryos (cont.) compared with embryos incubated from 24 to 48 hpf in PKF118-310 (PKF). Oral pole is at the top in all panels. All panels are the same magnification, scale bars 100 µm.

Following Wnt3-MO injection (Fig. 5A) or early treatments with PKF118-310 (Fig. 5B), the first phase of nematogenesis was disrupted, such that most Wnt3-MO embryos showed only a few nematoblasts clearly positive for Minicollagen3/4a. Mos3 expression was barely detectable and Znf845 cells reduced to trace levels or absent. This effect on nematoblast formation was reflected by reductions in the cell populations expressing Nanos1 and Piwi (Fig. 5; Supplementary File S3); by 48 hpf the Nanos1-expressing population was reduced to trace levels in many Wnt3-MO embryos, as expected given the continued expression of these mRNAs during early nematogenesis. In contrast we observed no obvious reductions in cell populations expressing Nanos1 (Fig. 5A,B) in Wnt3-MO or PKF118-310 treated embryos at 24 hpf, suggesting that initial i-cell formation and maintenance are not directly affected by Wnt signalling.

The early steps of development of neurons by both i-cell and ectodermal neurogenesis pathways were found to be sensitive to Wnt-β-catenin inhibition. Both Neurogenin and Hlh6 expression were strongly reduced in PKF118-310 treated embryos at 24 hpf, and ELAV expression was markedly reduced by both early and late PKF118-310 treatments. A further requirement for Wnt signalling in differentiation steps was shown by late treatments with PKF118-310 (Fig. 5C). This had no obvious effect on existing Znf845 or Minicollagen expressing cell populations, but blocked their differentiation into nematocytes: Nematocilin expression was detected only weakly, and no nematocytes were found in the ectoderm. Furthermore, after these late PKF118-310 treatments, the number of cells expressing ELAV was greatly reduced, indicating that neuronal differentiation is not maintained in the absence of Wnt signalling. Consistently, both PP5/RFamide and PP2/GLWamide populations were reduced to trace levels in most embryos following PKF118-310 treatment from 24 hpf. In contrast, in Wnt3-MO planulae at 48h, we detected no RFamide cells but did detect PP2/GLWamide cells despite the strong aboralised phenotype (Fig 5A). One possible explanation for this apparent discrepancy is that the planula aboral territory requires low levels of Wnt-β-catenin signalling for its development, as shown in *Nematostella* (Leclère et al., 2016). Given that several Wnt ligands are expressed in the *Clytia* planula in overlapping domains (Momose et al., 2008), we suggest that low level Wnt-β-catenin signalling may operate in Wnt3-MO treated planulae. These levels would mimic normal aboral conditions and promote GLWamide neuron development, whereas PKF118-310 would completely abolish both.

Taken together these results indicate that programmes of neurogenesis, including nematogenesis, both ectodermal and i-cell derived, are sensitive to Wnt signalling. Wnt signalling promotes both the formation of nematoblasts and their differentiation into nematocytes, and likewise early neurogenic processes as well as terminal differentiation, particularly in oral and trunk regions of the developing planula.

## Discussion

*Clytia hemisphaerica* offers an excellent opportunity to address the embryological origin of neural cell types during hydrozoan larval development. By combining gene expression profiling, bisection experiments and lineage tracing, we have uncovered two parallel pathways, schematized in Figure 6. At the onset of gastrulation, Piwi/Nanos1 positive cells arising in the oral pole presumptive endoderm domain of the early gastrula rapidly embark on a programme of nematogenesis. This pathway, like all the neurogenesis steps we examined, is strongly affected by disruption of Wnt-β-catenin signalling, explaining the previously observed strong downregulation of Znf845 and Mos3, which we now know are early nematogenesis markers (Chari et al., 2021), in Wnt3-MO embryos (Lapébie et al., 2014).

**Figure 6.**
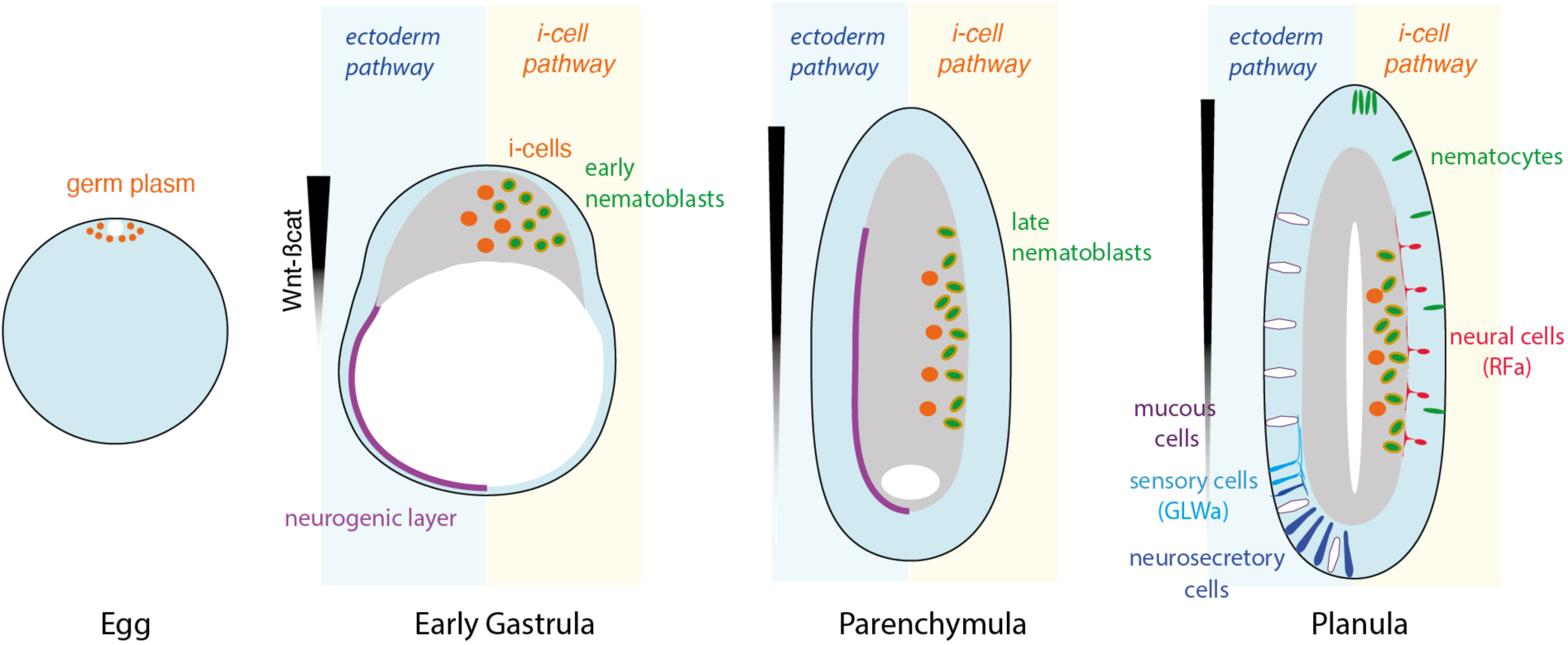
Dual neurogenesis pathways during *Clytia* larval development. Cartoon illustrating successive developmental stages from egg to planula. At the early gastrula, parenchymula and planula stages, cell types specific to the ectodermal pathway and i-cell pathway are depicted on the left and right sides respectively. See text for detailed discussion.

During normal planula development a large pool of Minicollagen3/4a-expressing nematoblasts accumulates in the gastrodermal region as early as the parenchymula stage. Only a small subpopulation of these subsequently migrate into the ectoderm and differentiate into mature nematocytes. We speculate that most of the ‘stalled’ nematoblasts will undergo rapid differentiation during metamorphosis to provide nematocytes for the primary polyp, enabling it to catch its first prey, as has been proposed for other hydrozoan planulae (Burmistrova et al., 2018; Chrysostomou et al., 2022). I-cells also generate some other neural cell types including RFamide expressing ganglionic cells.

In parallel to this i-cell pathway, a distinct neurogenesis process occurs in a basal layer of the aboral/lateral gastrula ectoderm generating GLWamide and possibly other neurosensory cells. Mucous cells and aboral secretory cells (Bodo and Bouillon, 1968) also derive from this layer. These two secretory cell types share molecular characteristics with neurons present in the planula and medusa (Ferraioli et al., 2026), and can be considered as part of a wider group of neurosecretory cell types or ‘paraneurons’, possibly with a common origin (Arendt et al., 2015; Babonis, 2025; Hobert, 2025).

### I-cell formation and fate during *Clytia* embryogenesis

The work reported here allows us to reassess the fate during *Clytia* larval development of i-cells, defined here as cells expressing genes typical of germ cell/stem cell pluripotency programmes (Juliano et al., 2010) such as Piwi, Vasa and Nanos1. Previously we showed that maternal “germ plasm” like granules including mRNAs of several of these genes is positioned around the female pronucleus at the egg animal pole (Leclère et al., 2012) as illustrated in orange in Figure 6, but found that Piwi/Nanos1 expressing i-cells could develop within the gastroderm from embryo fragments irrespective of whether or not they contained this germ plasm. We now know that most of the “i-cells” of the developing planula form de novo at the oral pole from the early gastrula stage, and that from this early stage they have already embarked upon nematogenesis as indicated by expression of Znf845, Mos3 and later Minicollagen (this study; Chari et al., 2021). This validates previous histological descriptions noting the appearance of distinguishable nematoblasts during gastrulation, well ahead of the appearance of any cells with morphologies typical of totipotent i-cells (Bodo and Bouillon, 1968). The previous report that Piwi/Nanos1 positive cells can develop in larvae derived from vegetal as well as animal fragments of early embryos (Leclère et al., 2012) can, in this new light, be accounted for by the onset of nematogenesis in these embryos, and does not necessarily represent de novo formation of totipotent i-cells. Some i-cells also generate RFamide neurons and possibly other neural cell types. Expression of various potential molecular regulators of neural development have been reported in cells both scattered within the endodermal region and in the developing aboral ectoderm (this study; Jager et al., 2011; Kraus et al., 2020; Lapébie et al., 2014). Consistently, shRNA-mediated downregulation of a SoxB gene, expressed in an i-cell type pattern in the gastrodermal region during *Hydractinia* planula development, prevented formation of RFamide expressing cells (Chrysostomou et al., 2022; Flici et al., 2017).

Whether pluripotent i-cells, not yet engaged in neurogenesis or nematogenesis, and/or cells committed to a germline fate, are present in *Clytia* embryos and larvae remains an important open question. Resolving this issue will require the identification of specific markers for pluripotent and germ line cells in *Clytia*, something that has so far remained elusive.

### Alternative neurogenesis pathways in hydrozoans

Our bisection and Dendra2 tracing experiments in *Clytia* showed that several neural cell types of the planula larva, including GLWamide cells, mucous cells and secretory cells, derive from the ectoderm. This was also the conclusion from previous studies based on EM of hydrozoan planulae manipulated in various ways to eliminate i-cells (Martin and Thomas, 1981; Thomas et al., 1987). This ectodermal neurogenesis pathway is characterised by expression in a basal, neurogenic, layer of the blastula and gastrula ectoderm of the transcription factor genes Sox10 (Kraus et al., 2020) and then Hlh6 expression (this study). Also expressed in this layer are a number of genes from families implicated in modulating neurogenesis in other species such as bZip as well as Wnt and Notch pathway regulators, identified by their upregulation in Wnt3-MO early gastrula-stage embryos (Lapébie et al., 2014). Strong expression of ELAV in this layer, even before gastrulation is complete, emphasises the important contribution of ectodermal neurogenesis during planula formation. This picture of embryo neurogenesis differs from that characterised in the adult stages, i.e. polyps, of *Hydractinia* (Song et al., 2025) and *Hydra,* where i-cells give rise to all neurons including GLWamide positive sensory cells (Mitgutsch, 1999; Siebert et al., 2008). scRNAseq data suggest that i-cells generate a similar range of neural cell types in *Clytia* medusa stages (Chari et al., 2021).

Ectodermal neurogenesis is typical of embryogenesis in bilaterian (protostome and deuterostome) animals and has also been described in anthozoan cnidarians such as *Nematostella,* where both sensory and ganglionic cells arise in the ectodermal epithelium (Lemaître et al., 2023; Nakanishi et al., 2012; Rentzsch et al., 2017), and in the scyphozoan *Aurelia* (Nakanishi et al., 2008). In *Nematostella,* neurogenesis in the gastroderm follows that in the ectoderm at the planula stage (Nakanishi et al., 2012). Furthermore, juvenile and adult gastrodermal tissues harbour Vasa2/Piwi1 expressing cells with similarities to hydrozoan i-cells, generating at least some of the neuronal precursors in addition to germ cells (Miramón-Puértolas et al., 2024). In hydrozoans, endodermal positioning of i-cells appears to be a particularity of the planula stage (Bodo and Bouillon, 1968; Leclère et al., 2012; Martin, 1991; Rebscher et al., 2008), perhaps relating to a requirement to stockpile nematoblasts for rapid deployment in the primary polyp following metamorphosis (see above). At other life cycle stages, i-cells are mainly found between cells at the base of the ectoderm, with the notable exception of the gonad at the medusa stage (Bosch et al., 2010; Hou et al., 2021; Jessus et al., 2020).

More generally, our study reinforces the hypothesis that neurogenesis from the ectodermal epithelium evolved in a common ancestor of Cnidaria and Bilateria (Nakanishi et al., 2012). Ectodermal neurogenesis would have persisted during evolution of Hydrozoa, at least during development of the planula larva. Under this scenario, i-cells may have emerged in Hydrozoa as a complementary mechanism for neurogenesis/nematogenesis. In the *Clytia* larva, i-cells give rise mainly to nematocytes but also at least some of the RFamide cells, while in the adult medusa nematocytes and most neurons derive from i-cells (Chari et al., 2021; Denker et al., 2008). We can thus speculate the marked stem-like capacity of hydrozoan i-cells to proliferate locally may have facilitated continuous rapid production of nematocytes at particular sites in adult hydrozoans.

## Acknowledgements

We thank D. Houston (Univ. Iowa) for his participation in pilot embryo cutting experiments, T. Lamonerie (Univ. Côte d’Azur) for participation in the Dendra2 experiments, our research group colleagues, C. Hudson and R. Copley for useful discussions, G. Jékely for generously donating the anti-RFamide antibody and O. Koizumi for the anti-LWamide antibody. Microscopy facilities were provided by the Imaging Platform and *Clytia* by the Service Aquariologie of Centre de Ressources Biologiques Marines (IMEV – FR 3761) of Institut de la Mer de Villefranche, both supported by EMBRC-France (ANR-10-INBS-02). The work was funded by the H2020/Marie Skłodowska-Curie ITN “EvoCell” (grant agreement no. 766053) to E.H and CNRS core funding to LBDV.

## List of SUPPLEMENTARY FILES

**S1) Gene names and accession numbers**

Details of the genes followed in this study.

**S2) Polarity maintenance in PKF treated larvae**

In situ hybridisation images of planula larvae (48 hpf) with probes to detect the genes indicated to the left of each panel, as markers for oral (Wnt3, Bra1) and Aboral (WgA1, FoxQ2a) polarity. Left column: Untreated planulae (cont.). Right column: incubated from 24-48 hpf in PKF118-310 (PKF). Oral pole is at the top in all panels. All panels are the same magnification, scale bars 100 µm.

**S3) Regulation of nematogenesis and neurogenesis by Wnt-β-catenin signalling**

Numbers of each type of in situ hybridisation patterns obtained with each probe. Yellow/ orange cells indicate the main phenotype for each group, as illustrated in Figure 5. Counts are from visual scoring of each stained planula in the corresponding group from one or two (*) experiment(s): ‘positive’ = clearly stained cells detected; ‘trace’ = staining observed but indistinct and/or very few cells. ND = not done.

