## Supplementary file S1 for "Dual origins for neural cells during development of the *Clytia* planula larva"

| Gene name | XLOC identifier | GenBank ID | Reference |
|---|---|---|---|
| | see Chari et al., 2021 Sci Adv | individual submissions | |
| piwi | XLOC_007915 | EU199802 | Leclère et al., 2012 Dev Biol |
| nanos1 | XLOC_036006 | AFD28590 | Leclère et al., 2012 Dev Biol |
| vasa | XLOC_033801 | JQ397273 | Leclère et al., 2012 Dev Biol |
| znf845 | XLOC_017841 | JAC85098 | Lapébie et al., 2014 PLoS Genetics |
| mos3 | XLOC_015554 | JAC85029 | Lapébie et al., 2014 PLoS Genetics |
| minicollagen 3/4a | XLOC_044122 | ABW02882 | Denker et al., 2008 Dev Biol |
| M14 peptidase | XLOC_039385 | | Chari et al., 2021 Sci Adv |
| nematocilin | XLOC_004102 | | Chari et al., 2021 Sci Adv |
| ELAV | XLOC_030971 | | Chari et al., 2021 Sci Adv |
| pp5 | XLOC_019434 | KX496951 | Takeda et al. 2018 Dev; Sinigaglia et al. 2018 Dev Biol |
| hlh6 | XLOC_030920 | KT318157 | this study |
| pp2 | XLOC_017096 | KX496948 | Takeda et al. 2018 Devt; Chari et al. 2021 Sci Adv |
| neurogenin | XLOC_018937 | KT318150 | this study |
