## Supplementary figures and images for "Dual origins for neural cells during development of the *Clytia* planula larva"

### Supplementary file S2

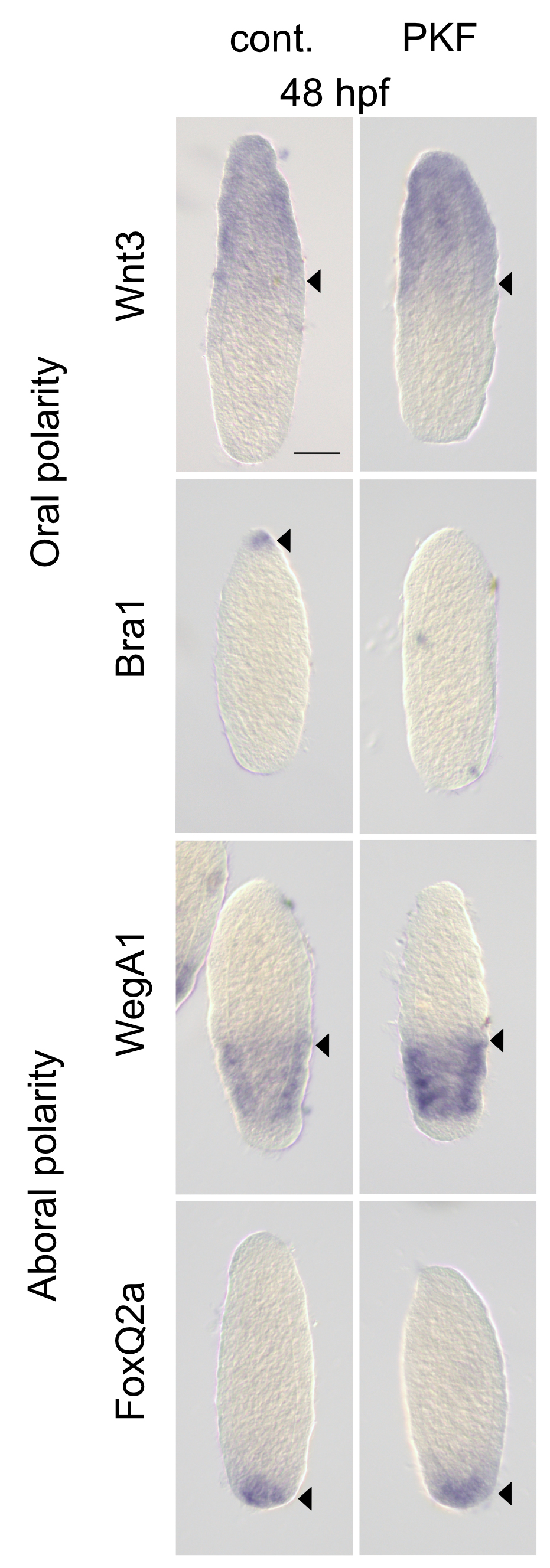
