## Supplementary file S3 for "Dual origins for neural cells during development of the *Clytia* planula larva"

**Supplementary File S3. Effects of Wnt inhibition on cell type development assessed by marker gene ISH**

| Probe | | control P24h | | | | Wnt3-MO P24h | | | | control P48h | | | | Wnt3-MO 48h | | | |
|---|---|---|---|---|---|---|---|---|---|---|---|---|---|---|---|---|---|
| | | positive | trace | none | total | positive | trace | none | total | positive | trace | none | total | positive | trace | none | total |
| Nanos1 | numbers | 10 | 3 | 0 | 13 | 13 | 2 | 1 | 16 | 6 | 1 | 0 | 7 | 7 | 12 | 0 | 19 |
| | % | **77** | **23** | **0** | | **81** | **13** | **6** | | **86** | **14** | **0** | | **37** | **63** | **0** | |
| Piwi | numbers | | ND | | | | ND | | | | ND | | | | ND | | |
| | % | | | | | | | | | | | | | | | | |
| Znf845 | numbers | 8 | 0 | 0 | 8 | 0 | 13 | 9 | 22 | 8 | 0 | 9 | 17 | 0 | 13 | 9 | 22 |
| | % | **100** | **0** | **0** | | **0** | **59** | **41** | | **47** | **0** | **53** | | **0** | **59** | **41** | |
| | numbers | 8 | 0 | 0 | 8 | 7 | 9 | 5 | 21 | 16 | 4 | 1 | 21 | 10 | 8 | 0 | 18 |
| | % | **100** | **0** | **0** | | **33** | **43** | **24** | | **76** | **19** | **5** | | **56** | **44** | **0** | |
| Mos3 | numbers | 24 | 0 | 0 | 24 | 5 | 17 | 7 | 29 | 38 | 0 | 0 | 38 | 2 | 19 | 18 | 39 |
| | % | **100** | **0** | **0** | | **17** | **59** | **24** | | **100** | **0** | **0** | | **5** | **49** | **46** | |
| | numbers | 10 | 0 | 0 | 10 | 0 | 5 | 12 | 17 | 5 | 5 | 0 | 10 | 0 | 10 | 5 | 15 |
| | % | **100** | **0** | **0** | | **0** | **29** | **71** | | **50** | **50** | **0** | | **0** | **67** | **33** | |
| minicol3/4 | numbers | 5 | 0 | 0 | 5 | 1 | 0 | 8 | 9 | 7 | 0 | 0 | 7 | 4 | 16 | 3 | 23 |
| | % | **100** | **0** | **0** | | **11** | **0** | **89** | | **100** | **0** | **0** | | **17** | **70** | **13** | |
| | numbers | | | | | | | | | | | | | | | | |
| | % | | | | | | | | | | | | | | | | |
| Nematocillin | numbers | | ND | | | | ND | | | 24 | 2 | 0 | 26 | 0 | 6 | 5 | 11 |
| | % | | | | | | | | | **92** | **8** | **0** | | **0** | **55** | **45** | |
| Neurogenin | numbers | | ND | | | | ND | | | | ND | | | | ND | | |
| | % | | | | | | | | | | | | | | | | |
| | numbers | | | | | | | | | | | | | | | | |
| | % | | | | | | | | | | | | | | | | |
| Hlh6 | numbers | | ND | | | | ND | | | | ND | | | | ND | | |
| | % | | | | | | | | | | | | | | | | |
| ELAV | numbers | | ND | | | | ND | | | | ND | | | | ND | | |
| | % | | | | | | | | | | | | | | | | |
| PP5 | numbers | | ND | | | | ND | | | 7 | 0 | 0 | 7 | 0 | 1 | 5 | 6 |
| | % | | | | | | | | | **100** | **0** | **0** | | **0** | **17** | **83** | |
| PP2 | numbers | | ND | | | | ND | | | 9 | 0 | 0 | 9 | 18 | 0 | 2 | 20 |
| | % | | | | | | | | | **100** | **0** | **0** | | **90** | **0** | **10** | |

| Probe | | control 24h | | | | PKF 24h | | | | control 48h | | | | PKF 48h | | | |
|---|---|---|---|---|---|---|---|---|---|---|---|---|---|---|---|---|---|
| | | positive | trace | none | total | positive | trace | none | total | positive | trace | none | total | positive | trace | none | total |
| Nanos1 | numbers | 17 | 0 | 0 | 17 | 14 | 0 | 0 | 14 | 40 | 35 | 15 | 90 | 12 | 12 | 10 | 34 |
| | % | **100** | **0** | **0** | | **100** | **0** | **0** | | **44** | **39** | **17** | | **35** | **35** | **29** | |
| Piwi | numbers | 26 | 0 | 0 | 26 | 21 | 0 | 0 | 21 | | | | | | | | |
| | % | **100** | **0** | **0** | | **100** | **0** | **0** | | | | | | | | | |
| Znf845 | numbers | 18 | 0 | 1 | 19 | 0 | 12 | 0 | 12 | 22 | 0 | 0 | 22 | 14 | 7 | 0 | 21 |
| | % | **95** | **0** | **5** | | **0** | **100** | **0** | | **100** | **0** | **0** | | **67** | **33** | **0** | |
| | numbers | 23 | 0 | 0 | 23 | 0 | 0 | 11 | 11 | | | | | | | | |
| | % | **100** | **0** | **0** | | **0** | **0** | **100** | | | | | | | | | |
| Mos3 | numbers | 36 | 0 | 0 | 36 | 9 | 4 | 0 | 13 | | ND | | | | ND | | |
| | % | **100** | **0** | **0** | | **69** | **31** | **0** | | | | | | | | | |
| | numbers | 14 | 0 | 0 | 14 | 9 | 0 | 3 | 12 | | | | | | | | |
| | % | **100** | **0** | **0** | | **75** | **0** | **25** | | | | | | | | | |
| minicol3/4 | numbers | 16 | 0 | 0 | 16 | 0 | 6 | 5 | 11 | 29 | 5 | 1 | 35 | 20 | 4 | 0 | 24 |
| | % | **100** | **0** | **0** | | **0** | **55** | **45** | | **83** | **14** | **3** | | **83** | **17** | **0** | |
| | numbers | 30 | 0 | 0 | 30 | 0 | 0 | 7 | 7 | | | | | | | | |
| | % | **100** | **0** | **0** | | **0** | **0** | **100** | | | | | | | | | |
| Nematocillin | numbers | | ND | | | | ND | | | 34 | 0 | 0 | 34 | 0 | 51 | 16 | 67 |
| | % | | | | | | | | | **100** | **0** | **0** | | **0** | **76** | **24** | |
| Neurogenin | numbers | 0 | 14 | 3 | 17 | 0 | 1 | 17 | 18 | | ND | | | | ND | | |
| | % | **0** | **82** | **18** | | **0** | **6** | **94** | | | | | | | | | |
| | numbers | 35 | 0 | 0 | 35 | 0 | 0 | 13 | 13 | | | | | | | | |
| | % | **100** | **0** | **0** | | **0** | **0** | **100** | | | | | | | | | |
| Hlh6 | numbers | 24 | 0 | 0 | 24 | 0 | 12 | 3 | 15 | | ND | | | | ND | | |
| | % | **100** | **0** | **0** | | **0** | **80** | **20** | | | | | | | | | |
| ELAV | numbers | 16 | 0 | 0 | 16 | 0 | 0 | 12 | 12 | 35 | 0 | 2 | 37 | 0 | 30 | 0 | 30 |
| | % | **100** | **0** | **0** | | **0** | **0** | **100** | | **95** | **0** | **5** | | **0** | **100** | **0** | |
| PP5 | numbers | | ND | | | | ND | | | 16 | 10 | 0 | 26 | 2 | 14 | 6 | 22 |
| | % | | | | | | | | | **62** | **38** | **0** | | **9** | **64** | **27** | |
| PP2 | numbers | | ND | | | | ND | | | 21 | 8 | 8 | 37 | 2 | 14 | 7 | 23 |
| | % | | | | | | | | | **57** | **22** | **22** | | **9** | **61** | **30** | |

Numbers are from visual scoring of each stained planula in the corresponding group of a single experiment: 'positive' = clearly stained cells detected; 'trace' = staining observed but indistinct and/or very few cells.
Yellow/orange cells indicate the main phenotype for each group, as illustrated in Figure 5.
